# *In situ* and high-resolution Cryo-EM structure of the Type VI secretion membrane complex

**DOI:** 10.1101/441683

**Authors:** Chiara Rapisarda, Yassine Cherrak, Romain Kooger, Victoria Schmidt, Riccardo Pellarin, Laureen Logger, Eric Cascales, Martin Pilhofer, Eric Durand, Rémi Fronzes

## Abstract

Bacteria have evolved macromolecular machineries that secrete effectors and toxins to survive and thrive in diverse environments. The type VI secretion system (T6SS) is a contractile machine that is related to *Myoviridae* phages. The T6SS is composed of a baseplate that contains a spike onto which an inner tube is built, surrounded by a contractile sheath. Unlike phages that are released to and act in the extracellular medium, the T6SS is an intracellular machine inserted in the bacterial membranes by a trans-envelope complex. This membrane complex (MC) comprises three proteins: TssJ, TssL and TssM. We previously reported the low-resolution negative stain electron microscopy structure of the enteroaggregative *Escherichia coli* MC and proposed a rotational 5-fold symmetry with a TssJ:TssL:TssM stoichiometry of 2:2:2. Here, cryo-electron tomography analysis of the T6SS MC confirmed the 5-fold symmetry *in situ* and identified the regions of the structure that insert into the bacterial membranes. A high resolution model obtained by single particle cryo-electron microscopy reveals its global architecture and highlights new features: five additional copies of TssJ, yielding a TssJ:TssL:TssM stoichiometry of 3:2:2, a 11-residue loop in TssM, protruding inside the lumen of the MC and constituting a functionally important periplasmic gate, and hinge regions. Based on these data, we revisit the model on the mechanism of action of the MC during T6SS assembly and function.

## Introduction

In a competitive environment, the ability to communicate and outcompete neighbours provides bacteria with key advantages to survive. The type VI secretion system (T6SS) is a macromolecular complex involved in the release of toxins that disrupt essential functions in competitor cells (Russell *et al*, 2014). The T6SS is associated with increased survival and pathogenicity in bacteria expressing it (Zhao *et al*, 2018). The T6SS is composed of 13-14 core proteins (Boyer *et al*, 2009), usually encoded in the same locus in the genome (Mougous *et al*, 2006). The T6SS assembles a molecular spring-loaded dagger, which punctures the target cell to secrete fully folded effector proteins into neighbouring bacteria (Russell *et al*, 2011) or eukaryotic hosts (Barker *et al*, 2009). The full assembly consists of the trans-envelope TssJLM membrane complex (MC) (Durand *et al*, 2015) that tethers the TssKFGE-VgrG baseplate (Brunet *et al*, 2015)(No Title), onto which the tail polymerize. This tail comprises the inner tube made of stacks of Hcp hexamers wrapped by the TssBC sheath proteins that polymerize in a helical conformation (Brunet *et al*, 2014; Clemens *et al*, 2015; Kudryashev *et al*, 2015; Chang *et al*, 2017) and tipped by the spike VgrG which can be sharpened by the PAAR protein (Renault *et al*, 2018; Shneider *et al*, 2013). Effectors are either associated within the Hcp lumen, or directly or indirectly bound to the VgrG or PAAR spike (Unterweger *et al*, 2017; Silverman *et al*, 2013; Shneider *et al*, 2013; Flaugnatti *et al*, 2016a; Quentin *et al*, 2018). Upon contact with a neighbouring cell, unknown signals trigger the contraction of the sheath causing the tube and spike to pierce the membranes and secrete the effectors (Basler *et al*, 2012).

While the baseplate, tube and sheath proteins are conserved among contractile injection systems, the MC is specific to the T6SS. TssJ is an outer membrane lipoprotein (Aschtgen *et al*, 2008a) that positions first at the site of assembly, and then recruits TssM and TssL (Durand *et al*, 2015). TssM and TssL are two inner membrane proteins that share homology with two accessory subunits associated with T4SSb, IcmF and IcmH/DotU (Aschtgen *et al*, 2012; Ma *et al*, 2009; Durand *et al*, 2012; Logger *et al*, 2016). Not only does the MC anchor the baseplate to the inner membrane, but it would also serves as a channel to allow the passage of the tail tube/spike and maintain the integrity of the attacking cell during the translocation of the inner tube (Durand *et al*, 2015). The different subunits and the MC have been extensively biochemically characterised and several crystal structures of the components are available from various bacterial species (Felisberto-Rodrigues *et al*, 2011; Durand *et al*, 2012; Rao *et al*, 2011; Robb *et al*, 2012; Chang & Kim, 2015; Durand *et al*, 2015). We previously reported the negative-stain electron microscopy (EM) structure of the TssJLM complex from enteroaggregative *Escherichia coli* (EAEC). We determined that 10 TssJ lipoproteins are bound to 10 TssM proteins, forming two concentric rings of pillars and arches that span the periplasm. The arches were shown to link to a flexible base composed of the N-terminal part of TssM and 10 copies of TssL (Durand *et al*, 2015). This study also revealed that the EAEC TssJLM complex assembles into a 5-fold rotationally symmetric trans-envelope structure. However, the symmetry mismatch between the 5-fold symmetry of the MC and 6-fold symmetry of the baseplate (Nazarov *et al*, 2018) questions whether the purified TssJLM MC reflects the *in vivo* situation. In addition, although most of the available crystal structures can be fitted into this EM structure, we currently lack molecular details on the whole complex, such as the precise location of the membranes, of the trans-membrane helices, and the potential presence of a periplasmic channel.

Here, we first present the *in situ* cryo-electron tomography (cryo-ET) structure of the EAEC TssJLM MC. These data confirm the 5-fold symmetry of the complex *in vivo*, and provide information on the location of the inner and outer membranes. We then present the single particle (SPA) cryo-electron microscopy (cryo-EM) structure of the MC that describes the intricate molecular architecture of the periplasmic portion of the complex. This high-resolution cryo-EM structure reveals the presence of five additional copies of TssJ at the tip of the complex, the presence of a periplasmic gate that closes a periplasmic channel. Finally, we demonstrate that the periplasmic gate and the additional TssJ are required for T6SS-dependent activity *in vivo*.

## Results

### Structure of the T6SS MC within the cell envelope

To observe the T6SS MC in its native cellular environment, we performed cryo-ET on bacterial cells (Weiss *et al*, 2017). With the aim to have a sufficient number of particles to obtain an *in situ* structure by subtomogram averaging, we imaged *E. coli* BL21(DE3) cells in which TssJLM were heterologously overexpressed. This strain does not possess T6SS genes, thereby preventing any crosstalk or protein-protein interactions between TssJLM and other natively present T6SS proteins. Since *E. coli* is too thick to be directly imaged by cryo-ET, we pursued three different approaches to tackle sample thickness. The first approach consisted of engineering a minicell-producing skinny strain (Farley et al, 2016) of *E. coli* BL21(DE3), and thereby generating a minicell strain that is compatible with the T7-based expression system. Although this strain produced minicells as small as 450 nm in diameter (Fig EV1A), their size still affected the contrast to an extent that did not allow for sub-tomogram averaging. Nevertheless, characteristic inverted Y-shaped particles (side views, Fig EV1B) and star-shaped particles (top views, Fig EV1C) could occasionally be observed in the periplasm of these minicells. In a second approach, tomograms were recorded of cells that were partially lysed and exhibited a high contrast (Fig EV1D, E). These cells, which have previously been described as “ghost cells” due to their transparent appearance, still had mostly intact membranes and conserved their cytoplasmic macromolecular complexes such as ribosomes (Fu *et al*, 2014). In a third approach, we used a state-of-the-art cryo-sample thinning method called focused ion beam (FIB) milling to thin *E. coli* BL21(DE3) cells in which TssJLM were heterologously overexpressed (Fig EV1F, G). FIB milling allows to etch through a lawn of bacterial cells and thin them down to under 200 nm (Medeiros *et al*, 2018a, 2018b; Marko *et al*, 2007). This approach was more native, as it was performed on intact rod-shaped wild-type BL21(DE3) cells. These tomograms had a high contrast and provided great detail. To confirm the relevance of these observations, we carried out control experiments in the native T6SS^+^ EAEC 17-2 strain. TssJLM particles were also readily observed by cryo-ET, both when heterologously overproduced (Fig EV1H, note that the particles could occasionally be found detached from the outer membrane: Fig EV1J) as well as under native conditions, *i.e.* in wild-type EAEC 17-2 cells (Fig EV1I).

Sub-volumes of star-shaped (top views) and inverted Y-shaped (side views) particles were manually picked and subsequently aligned and averaged. The resulting average was similar to the *in vitro* T6SS MC structure published previously (Durand *et al*, 2015), with a tip and a core, made of 5 pairs of pillars forming a narrow central channel, that splits into 10 arches (Fig 1A-E). Importantly, the 5-fold symmetry was evident without applying symmetry (Fig 1A’). As it is the case for the TssJLM MC solved by EM (Durand *et al*, 2015), the periplasmic core and the arches were well resolved, whereas the inner membrane embedded base and the outer membrane-embedded cap were poorly resolved. The Fourier shell correlation (FSC) curve indicated a resolution of 25 Å at a coefficient of 0.5 (Fig EV2 A). The prevalence of top views indicated that the average might be affected by a missing wedge (Fig EV2 B).

**Figure 1.**
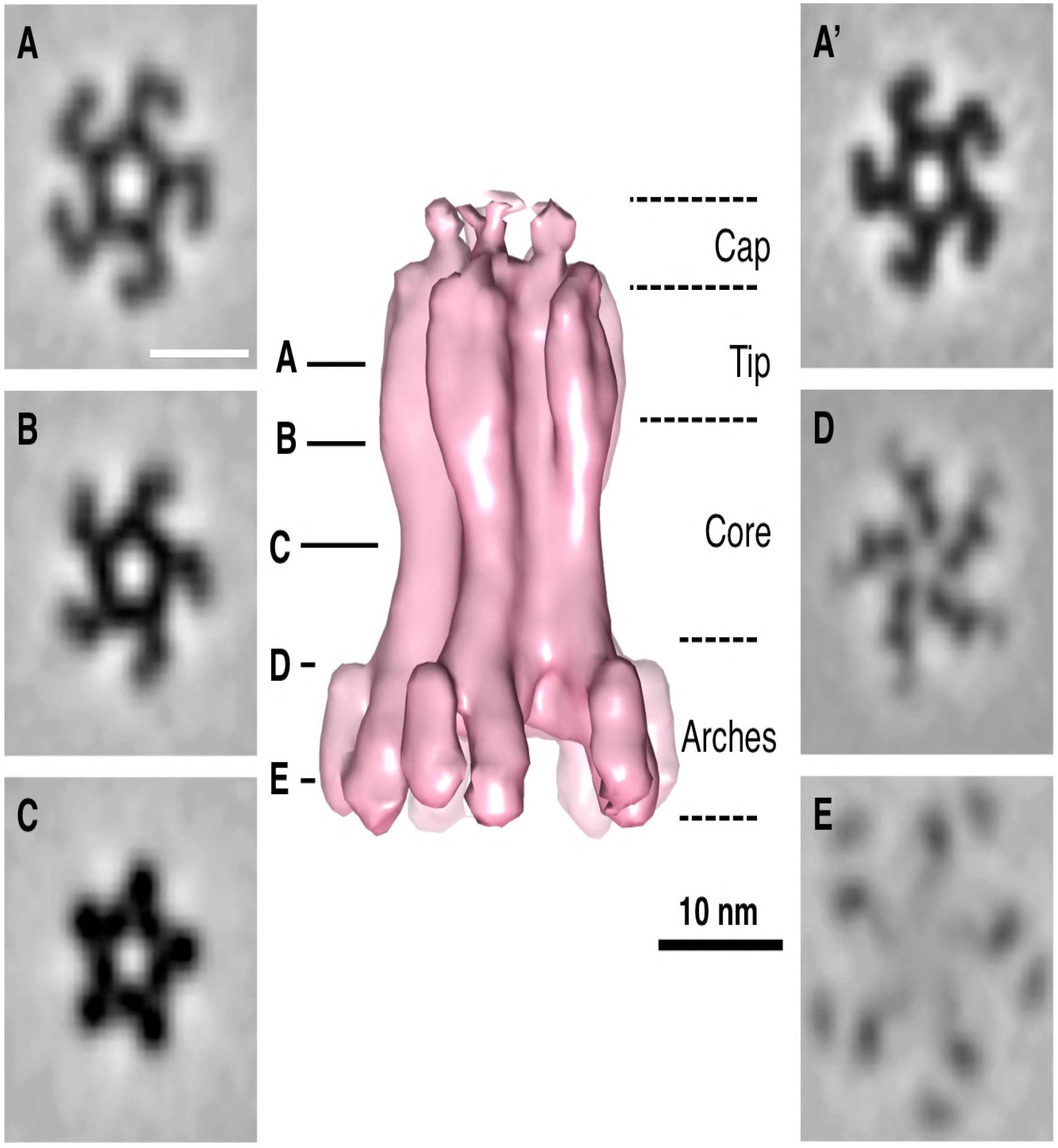
Subtomogram average of the TssJLM complex *in situ*. **A-E.** Isosurface of the subtomogram average (in pink) with an applied C5 symmetry and 0.69 nm tomographic slices (A-E) at the indicated heights. The average is shown in side view, whereas the slices represent perpendicular slices. The 5 pairs of pillars formed a narrow central channel and separated into 10 arches towards the base (base not visible at the used threshold). The division of the structure into subparts was adapted from (Durand *et al.*, 2015). Subvolumes were extracted from cryotomograms of ghost cells and FIB-milled intact *E. coli* BL21 cells heterologously expressing TssJLM. **A’**. Slice (0.69 nm) through the non-symmetrized average. Note that the C5 symmetry was visible.

After aligning and averaging a set of sub-volumes, it can be useful to place the isosurface of the average back in the original tomographic volume to analyze the positions and orientations of the individual aligned particles. In this way, we obtained a clear view of the location of the MC within the cell envelope (Fig 2A, Movie 1 and Fig EV2 C-E, Movie 2). The position of the membranes with respect to the TssJLM highlighted that the tip was embedded in the outer membrane without crossing it, while the MC was anchored in the inner membrane at the lower part of the arches (Fig 2B). In some cases, densities could be seen spanning the inner membrane and extending into the cytoplasm (Fig 2C). In what could be an overexpression artefact, TssJLM particles were also found in cytoplasmic membrane invaginations (Fig 2A). Altogether, these data confirmed the 5-fold symmetry of the T6SS TssJLM MC *in situ* and provided further insights into the position of the inner and outer membranes.

**Figure 2.**
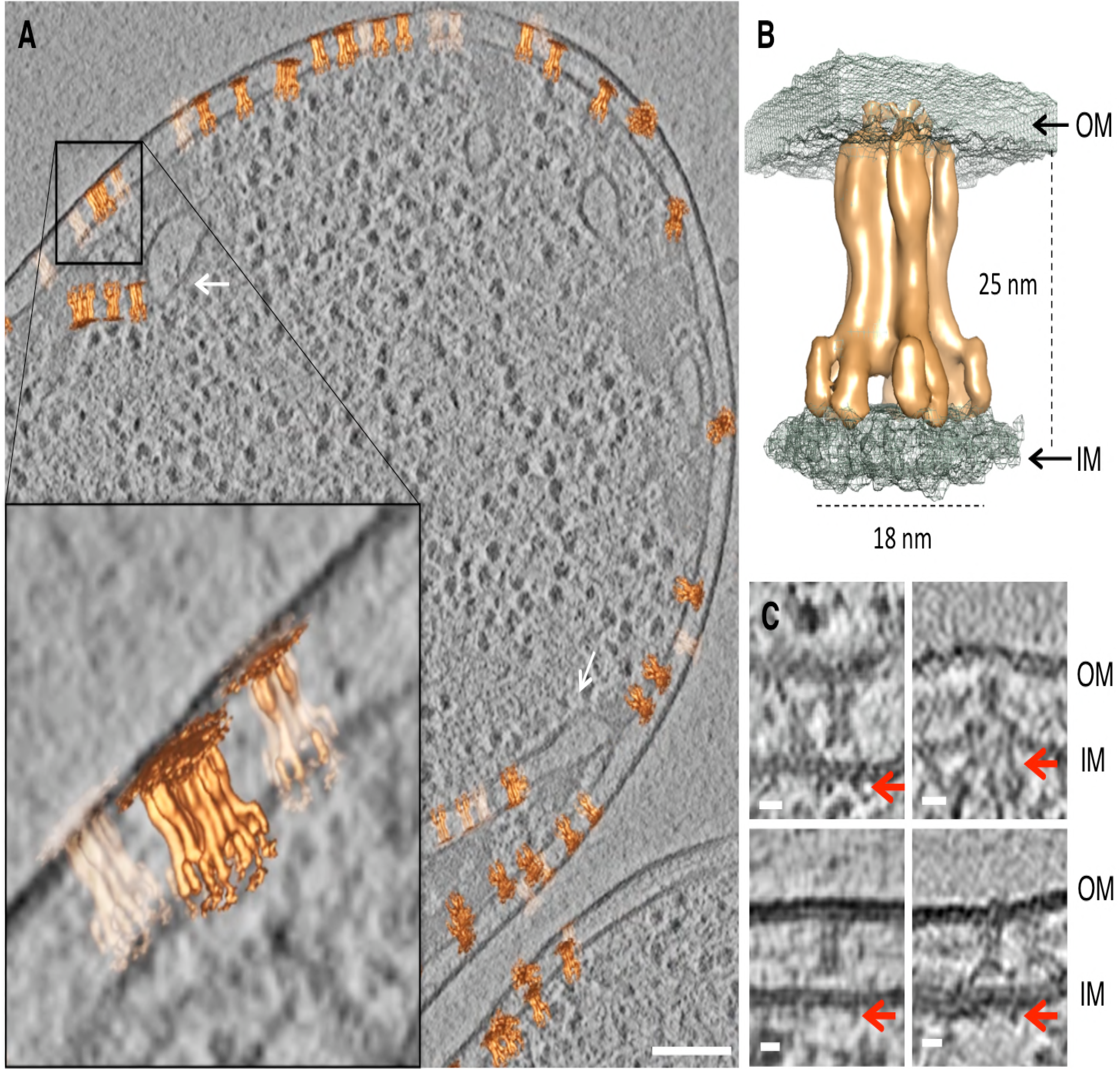
Position of TssJLM in the cell envelope. **A.** Slice (9.7 nm) through a cryotomogram of a FIB-milled *E. coli* BL21 cell expressing TssJLM. The average shown in Figure 1 was placed back at the positions and orientations of the individual subvolumes that were used to generate the final average. The zoomed-in area highlights the position of the TssJLM particles within the inner and the outer membrane. Some particles were found in cytoplasmic membrane invaginations, as indicated by white arrows. Scale bar 100 nm. **B.** Isosurface of the final average (orange) merged with the isosurface of a second average (with higher threshold; grey mesh). The panel shows the positioning of TssJLM with respect to both inner (IM) and outer membranes (OM). The distances corresponded to the widest and longest dimensions of the TssJLM complex. **C.** Cryotomographic slices (9.7 nm) showing side views of TssJLM. In these examples, the basal parts of TssJLM (red arrows) could be seen extending into the cytoplasm. Scale bar 10 nm.

### Structural analysis of the TssJLM complex by cryo-EM

We previously reported the negative stain EM structure of the EAEC TssJLM MC (Durand *et al*, 2015). Here, we used the same purification procedure and used SPA cryo-EM to obtain the cryo-EM structure of the 1.7 MDa T6SS MC at 4.9 Å overall resolution. With a 5-fold symmetry imposed, the local resolution ranged between 4.1 Å and 21.7 Å (Figs 3A and EV3A-E). The angular distribution of the particles was good and allowed us to reach a high resolution (Fig EV3 C, F).

**Figure 3.**
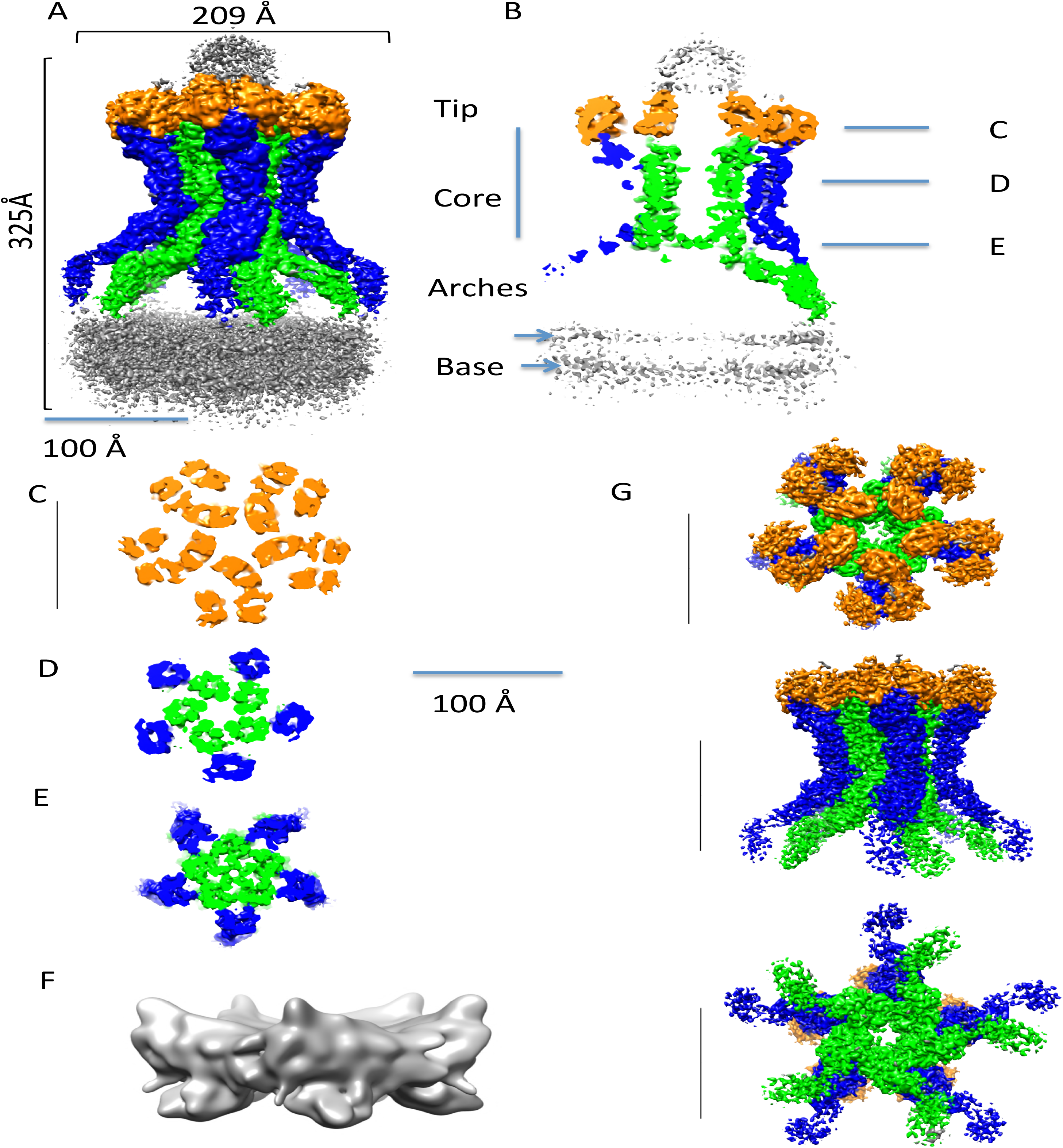
3D structure of the TE complex of the T6SS. **A.** Autosharpened Cryo-EM density of the full TE complex composed of TssJ, TssL and TssM. The inner pillars are coloured in green and the outer pillars in blue. The unstructured tip and base are in gray, while the top of the core is in orange. **B.** Vertical cross section of the cryo-EM density. Colouring is according to A. The position of the slices for C-E are indicated. The density of the base appears to be composed of two leaflets, indicated by 2 arrows. **C-E.** Cross sections of A, sliced at positions indicated in B. **F.** Subtracted, masked and unsharpened cryo-EM density of the mostly unstructured base **G.** Locally sharpened cryo-EM density of the subtracted and masked core. Three different views are shown. The colouring is according to A

The 5-fold symmetry is clearly visible in the 2D classes top views and the volume slices of the reconstruction, which retain a 5-pointed star shape (Fig EV3C,G). When no symmetry was applied during the reconstruction, the overall resolution decreased to 7.5 Å but the pentameric nature of the complex was maintained, with only one of the 10 arches displaying weaker density than the others causing the drop in resolution (Fig EV3H, I). This could indicate partial assembly of the complex *in vivo* or disassembly during its purification.

The overall structure resembles that obtained by negative stain (Fig EV3J(Durand *et al*, 2015)). The architecture of the complex comprises a tip connected to a core region that extends to a base, through a double ring of pillars with arches in proximity to the inner membrane (Fig 3A-B). Several notable features are already evident from the cryo-EM density map of the full complex (Fig 3A). First, the tip region and the base are disordered, and appear to be filled by random densities. Second, cross-sections of the full complex show that the channel, across which tube/spike transport might occur, is closed by a gate at the intersection between arches and pillars, above the inner membrane (Fig 3B-E).

To better characterise the flexibility of the base of the complex, we collected tomograms on the same frozen EM grids that were used to collect the SPA dataset (Fig EV4A-C). The base of individual complexes appeared as very heterogeneous, with single arches often pointing in opposite directions, or on the contrary several arches clumping together (Fig EV4A). On the other hand, the core was rigid and resembled a 5-branched star. Moreover, about 20% of the particles possessed only 3 or 4 out of 5 branches (Fig EV4B), indicating that partial assemblies could be stable. The particles were lying in a thin layer of ice (25 nm) and were found at different orientations (Fig EV4C1-2).

### The structure of the base

To try and overcome the inherent flexibility of the complex and better discern different features of the base, we performed a density subtraction of the tip, core and arches followed by a focused refinement of the base with and without symmetry applied. We thus obtained 2D classes and a 3D structure of the base at 17-Å resolution when a C5 symmetry was applied (Figs 3F and EV4E-F). When observing the cross section of the base in the full complex, as indicated by arrows in Fig3B, and in the subtracted structure (Fig EV4E), two 110 Å-wide linear densities are clearly visible, separated by 40 Å. This double layer of density is consistent with the density diagram of a lipid bilayer with the head groups being the most dense at a distance of ~4 nm from each other and fits well a lipid bilayer composed of PE, obtained using the CHARMM-GUI (Jo *et al*, 2007) (Fig EV4E). We conclude that the inner membrane sub-domain of the T6SS MC is filled by a lipid bilayer.

### An additional TssJ is present in the full complex

To focus on the best-resolved region of the cryo-EM map, the base was subtracted from the tip, a 2D classification and a masked 3D refinement was performed to obtain the structure of the core at 4.5 Å (Figs 3G and EV5A-B), with a local resolution ranging from 4 Å to >10 Å (Fig EV 5C). The known crystallographic structure of the C-terminus of TssM (aa. 869-1129) bound to TssJ (PDB 4Y7O) could be easily fitted in the outer and inner pillars, with a correlation of 0.8505 and 0.565 respectively, leaving an extra density (Fig EV5D-E). Interestingly, an extra TssJ subunit, TssJ’, which was not observed in the low resolution complex (Durand *et al*, 2015) fits in this extra density with a correlation of 0.879 (EV5E-F). We conclude that the T6SS MC comprises 15 TssJ proteins, 3 TssJs for 2 TssM (Fig 4A). No additional residues were visible for the N- and C-termini of TssJ (1-21 and 151-155), disordered in both the crystal and the cryo-EM structures. After placing the extra TssJ, we refined the TssM-TssJ crystal structure and the newly fitted TssJ copy against the cryo-EM map to obtain the structure of each TssM and TssJ monomer within the whole MC.

**Figure 4.**
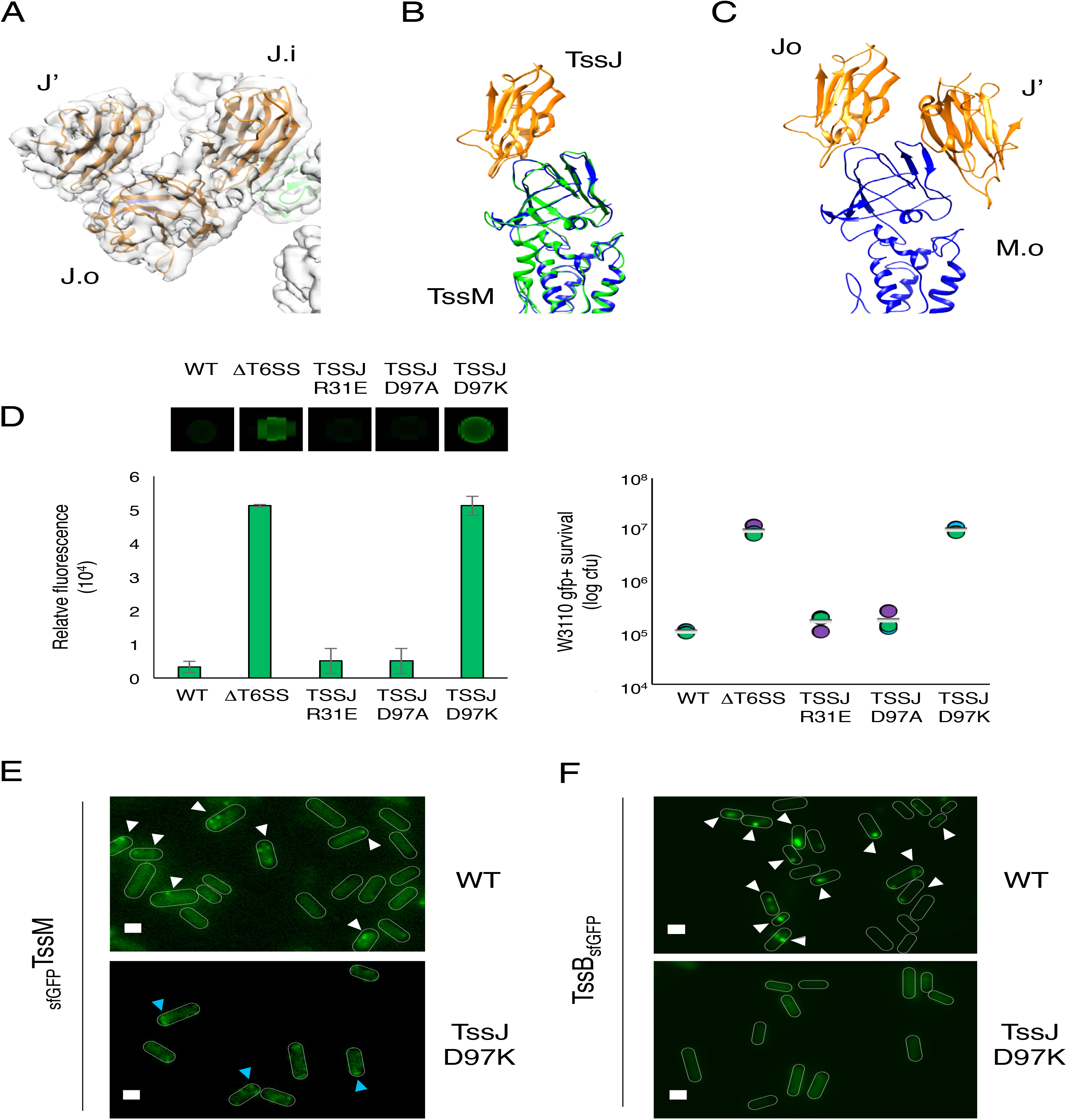
The TssJ’ monomer and its function. **A.** Ribbon diagram and locally sharpened surface representation (transparent= of the three TssJ protomers in orange, labelled TssJ.a, TssJ.A and TssJ’ for the internal, external and additional monomer respectively. In light green and blue are the TssM.a and TssM.A protomers respectively. **B.** Superimposition of the ribbon diagrams of TssM-TssJ heterodimer within the internal (green-orange) and the external (blue-orange) pillars. **C.** Ribbon diagram of the TssM.A (external pillar) with the TssJ.A and the TssJ’ (orange) **D.** Antibacterial assay. Prey cells (Gfp+ kanR E. coli W3110) were mixed with the indicated attacker cells, spotted onto Sci-1-inducing medium (SIM) agar plates and incubated for 4 h at 37°C. The image of a representative bacterial competition spot is shown on the upper part. The relative fluorescent level (in AU) and the number of recovered E. coli recipient cells are indicated in the lower graph (in log10 of colony forming units (cfu)). The assays were performed from at least three independent cultures, with technical triplicates and a representative technical triplicate is shown. The circles indicate values from technical triplicate, and the average is indicated by the bar. **E.** Fluorescence microscopy recordings showing sfGFPTssM foci in the parental (WT) and TssJ mutated strains (TssJ D97K). TssM foci containing cells are indicated by arrowheads. Microscopy analyses were performed independently three times, each in technical triplicate, and a representative experiment is shown. Scale bars, 1 μm. **F.** Fluorescence microscopy recordings showing TssBsfGFP sheath in the parental (WT) and TssJ mutated strains (TssJ D97K). Fluorescent sheath containing cells are indicated by arrowheads. Microscopy analyses were performed independently three times, each in technical triplicate, and a representative experiment is shown. Scale bars, 1 μm.

TssJ’ binds to the MC through previously unknown interfaces. If we consider the TssJ.i, TssJ.o and the TssJ’ subunits (Figs 4A-C and EV6A-B) with TssJ.i and TssJ.o being in contact with TssM.i and TssM.o respectively, they sit in the same position as in the crystal structure and in the outer and inner pillars (Fig 4B). By contrast, TssJ’ binds to TssM.o and TssJ.i strongly (Fig 4C, Table EV1). In particular, the interaction of TssJ’ with TssJ.i is specifically strong, as their contact is mediated not only by hydrogen bonds but also by salt bridges (R31 with E34, and D97 with R33; see Fig EV6C). TssJ’ binds to TssM.i via hydrogen bonds only and this interaction is comparable to that between TssJ.i and TssJ.o with TssM.i and TssM.o respectively (surface of interaction 573 Å^2^ and −2.6 kcal/mol ∆G) (Fig 4B-C, Table EV1).

### TssJ’ is required for MC assembly and T6SS activity

To gain further information on the function and *in vivo* relevance of TssJ’, mutations that specifically impact the TssJ’-TssJ.i interface were engineered onto the chromosome, at the native locus. Two residues, Arg-31 and Asp-97 (Fig EV6C), were targeted as they form a salt bridge with E34 and R33 in TssJ.a, respectively (Table EV 1 and Fig EV6C). The R31E, D97A and D97K substitutions were then tested for their ability to outcompete a fluorescent *E. coli* competitor strain. Although the R31E and D97A did not significantly impact T6SS antibacterial activity, the D97K mutation abolished proper function of the T6SS (Fig 4D). The assembly and stability of the T6SS MC was then assessed by fluorescence microscopy using a chromosomally-encoded and functional fusion protein between TssM and the super-folder GFP (_*sf*GFP_TssM, Durand *et al*, 2015) (Fig 4E). As previously observed (Durand *et al*, 2015), the _sfGFP_TssM strain forms stable foci. By contrast, cells producing the TssJ D97K variant form small and unstable fluorescent _sfGFP_TssM foci (Fig. 4E). Time-lapse fluorescence microscopy recordings showed that about 90% of the foci observed in the _sfGFP_TssM strain (n = 50) are stable over the 600-sec recording time, in agreement with the previous observation that the EAEC T6SS MC is stable and serves for several contraction/elongation cycles (Durand et al., 2015). By contrast, with a mean lifetime of ^~^ 107 sec (n = 50) the _sfGFP_TssM fluorescent foci observed in cells producing the TssJ D97K variant are drastically less stable (Fig EV 7A). We then tested whether the TssJ D97K variant promotes sheath assembly, using chromosomally-encoded functional _sfGFP_TssB fusion. Fig. 4F shows that, contrarily to the wild-type parental cells in which dynamic sheaths can be observed, no sheath polymerization occurs in presence of TssJ-D97K (Fig. 4F and EV Fig.7B). Altogether, these results demonstrate that TssJ’-TssJ.i interface is required for the stability of the T6SS MC, sheath formation and T6SS antibacterial activity.

### A flexible hinge within the TssM periplasmic domain

We were able to confidently build *de novo* the periplasmic domain of TssM including its N-terminal fragment (residues 579 to 869) that was missing in the crystal structure (Durand *et al*, 2015). This was done in Coot (Emsley *et al*, 2010) by using Phyre2 secondary structure predictions (Kelley *et al*, 2015) and RaptorX evolutionary covariance interaction predictions (Källberg *et al*, 2012) as validation tools (Fig 5A and EV8A-C). The cryo-EM structure of the TssM periplasmic domain slightly differs from the X-ray structure (Fig EV9A). From the C-terminus, helix 869-891 extends to amino-acid 841 with a slight kink at residue Pro-870. The remaining N-terminal fragment forms an α-helical domain comprising 8 helices that snake back and forth to the inner membrane (Fig 5A). The region closest to the membrane was too flexible to be resolved and for an atomic model to be built (Fig EV8B). The cryo-EM pseudoatomic model of the fully assembled TssM-TssJ complex shows it forms a bell shape composed of two rings of pillars that twist around each other (Fig 5A). Within each asymmetric unit, two copies of TssM are present, named TssM.o in the external pillar and TssM.i in the internal pillar. The inner and the outer pillar TssM proteins are superimposable with the exception of a 23° kink located at residue 867 (Fig EV9B). These two TssM subunits interact front-to-back (Fig EV9C) with an area of 1168 Å^2^, a binding energy of −9.9 kcal/mol, and a ∆G of −5.2 kcal/mol. Each TssM.i also interacts with two adjacent TssM.i within the inner TssM ring at an angle of 76°, with an area of 1529 Å^2^, a binding energy of −12 kcal/mol and a ∆G of −5.32 kcal/mol (Figs EV6A and EV9D). Finally, TssM.o^+1^ also interacts with TssM.i and the two are oriented at 68° from one another (Figs EV6A and EV9E).

**Figure 5.**
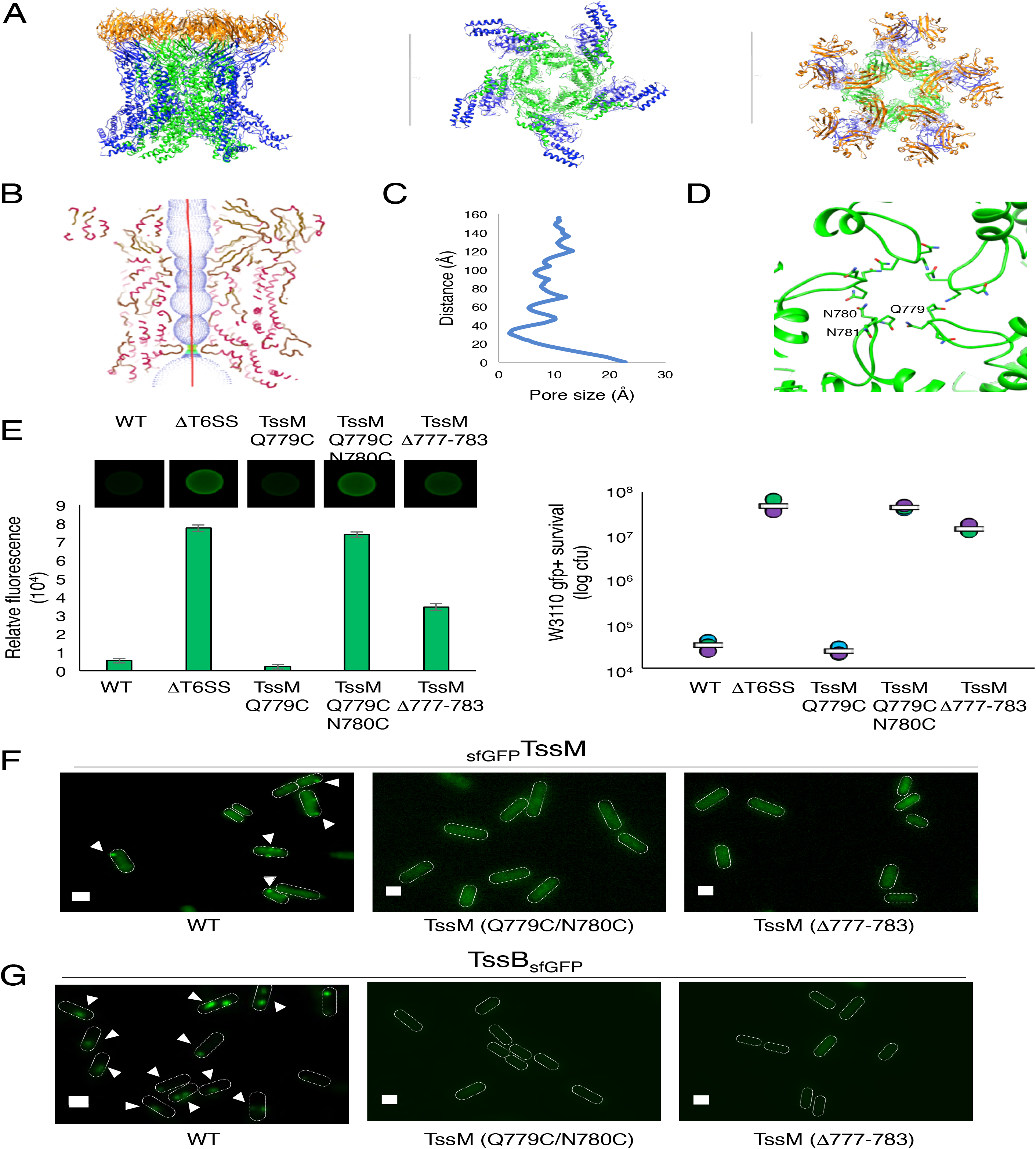
The TssM protein and the periplasmic gate. **A.** The high-resolution structure of MC complex in different orientations. TssJ is in orange, TssM is in green and blue according to their position as an internal or external pillar respectively. **B.** The pore radius formed by the internal pillar of TssM protomers is highlighted as dots and mapped using HOLE. The atomic model of TssM is coloured according to secondary structure (Helices in pink and strands in brown) **C.** A graph showing the pore radius calculated along the pore centre of the internal pillar of TssMs. **D.** Ribbon diagram of the region surrounding the periplasmic gate, with the amino acids involved in the formation of the gate in atom form. Gln779 and Asn780-781 from the internal pillars of TssM (green) are labelled in one of the protomers **E.** Antibacterial assay. Prey cells (Gfp+ kanR E. coli W3110) were mixed with the indicated attacker cells, spotted onto Sci-1-inducing medium (SIM) agar plates and incubated for 4 h at 37°C. The image of a representative bacterial competition spot is shown on the upper part. The relative fluorescent level (in AU) and the number of recovered E. coli recipient cells are indicated in the lower graph (in log10 of colony forming units (cfu)). The assays were performed from at least three independent cultures, with technical triplicates and a representative technical triplicate is shown. The circles indicate values from technical triplicate, and the average is indicated by the bar. **F.** Fluorescence microscopy recordings showing sfGFPTssM foci in the parental (WT) and TssM mutated strains (TssM Q779C/N780C, TssM ∆777-783). TssM foci containing cells are indicated by arrowheads. Microscopy analyses were performed independently three times, each in technical triplicate, and a representative experiment is shown. Scale bars, 1 μm. **G.** Fluorescence microscopy recordings showing TssBsfGFP sheath in the parental (WT) and TssM mutated strains (TssM Q779C/N780C, TssM ∆777-783). Fluorescent sheath containing cells are indicated by arrowheads. Microscopy analyses were performed independently three times, each in technical triplicate, and a representative experiment is shown. Scale bars, 1 μm.

A poorly-defined density that sits in the core region between TssM.i and TssM.o^+1^, was attributed to the C-terminus of TssM. We built a small loop that terminates into a helix. (Fig EV9F). This region is disordered in the outer pillar monomer (TssM.o). Additionally, the resolution of the pillars gets worse towards the basal side, and no secondary structure could be identified when we tried to build *de novo* the atomic model of TssM. Despite this high degree of flexibility, we produced a model of the periplasmic region between amino acids 390 and 550 based on a RaptorX contact predictions. This model fits well (correlations of 0.819 and 0.827) into the remaining densities (Fig EV9G). The EM density shows that while they are separated at the level of the arches, the inner and the outer pillar re-join at the level of the inner membrane (Fig EV9G).

### The periplasmic TssM gate

The inner pillars of TssM form a channel with a diameter that varies between 2.6 Å and > 20 Å (Fig 5B-C). The site of constriction observed in the cryo-EM density (Figs 3B,E) corresponds to loop 776-786 in the atomic model of TssM (Fig 5D). Specifically, residues Gln-779 and Asn-780-781 maintain the constriction via polar interactions (Fig 5D). In the external pillar, the same loop interacts with loop 600-625 on the neighbouring pillar, providing further stabilisation of the structure (Fig EV10A). Conservation analysis of the sequence with related proteins predicted by the ConSurf server (Celniker *et al*, 2013), indicated that the sequence of the loop 776-786 is poorly conserved amongst species (Fig EV10B) although the presence of a loop at this position is a conserved feature. As previously proposed (Durand *et al*, 2015), these data suggest that the purified TssJLM MC is in a closed state. Large conformational changes, including modification of the constriction and movement of the inner pillars are therefore required to allow the passage of the tube/spike during sheath contraction.

### The TssM periplasmic gate is required for MC assembly and T6SS function

To test the function of the periplasmic gate, several mutations were engineered at the *tssM* locus in the wild-type EAEC 17-2 strain and the function of the T6SS was assessed as previously described (Fig 5E-G). To covalently stabilize the contacts between the internal pillars and thus prevent MC opening, Gln779 and Asn780 were substituted with cysteines (Q779C-N780C). Conversely, a constitutively open gate was created by deleting a large portion of the constriction loop (∆777-783). Antibacterial competition assays showed that the Q779C-N780C variant loses the ability to outcompete competitor cells, whereas the single control mutant Q779C did not (Figure 5E). Deletion of the ∆777-783 loop also impaired the T6SS antibacterial activity (Figure 5E). Fluorescence microscopy recordings further show that _sfGFP_TssM Q779C-N780C and _sfGFP_TssM ∆777-783 do not assemble TssM foci since, in contrast to the parental strain, diffuse fluorescent pattern were observed (Figs 5F and EV11). These results demonstrate that the MC is not properly assembled when the integrity of the periplasmic gate is impacted. As expected, these mutant cells were not able to assemble T6SS sheaths (Figs 5G and EV11).

## Discussion

In this study, we report the *in situ* and *in vitro* structures of the T6SS TssJLM MC from EAEC. The cryo-ET structure confirmed the 5-fold symmetry and general architecture *in vivo* while the high-resolution cryo-EM structure provided molecular details about the periplasmic portion of the complex (Fig 1, Fig 3 and Movie 3). As previously defined, the pentameric propeller-like structure composed of 10 pillars intertwined with each other was observed both *in situ* and from purified material (Figs 1 and 3C-E). Cryo-ET analyses allowed to position both the inner and outer membranes. As anticipated based on biochemical experiments showing that TssJ is a periplasmic lipoprotein attached to the outer membrane by an acyl anchor (Aschtgen *et al*, 2008), the tip of the complex is embedded in the outer membrane (Fig 2B). However, neither the predicted detergent cap in the SPA cryo-EM structure nor the cryo-ET data, indicate that TssJLM breaches or crosses the outer membrane (Fig 2, Fig 3A-B). Nevertheless, it has been shown that this region is extracellularly exposed in wild-type EAEC cells (Durand *et al*, 2015). Although the cryo-EM structure demonstrates that the C-terminal region of TssM locates in the periplasm, we propose that the recruitment of specific T6SS components induces MC conformational changes and cell surface exposition of the TssM C-terminus. While our previous study suggested that the inner membrane locates at the level of the arches (Durand *et al*, 2015), the cryo-ET analyses revealed that the inner membrane surrounds the base (Fig 2B). Moreover, some tomograms also revealed the most basal parts of the MC, which cross the inner membrane into the cytoplasm (Fig 2A, C). The *in situ* cryo-ET structure presents an apparent elongation of the tip region compared to the cryo-EM structure (Fig EV12A, B and Movie 3), which could correspond to an additional density associated to the outer membrane or which could alternatively be explained by the missing wedge. Nevertheless, the similarities between both structures allowed the atomic model of TssJ - TssM to be docked into the *in situ* average (Fig EV12C, D), whereas the cryo-EM structure could be placed in a cellular context (Fig EV12E and Movie 3).

The base of the complex in this higher resolution structure was not better resolved than in the negative stain structure (Figs 3A, F, and EV3D). This base should comprise 10 copies of the TssM and TssL cytoplasmic domains (Durand *et al*, 2015). TssL forms dimers (Zoued *et al*, 2018; Durand *et al*, 2012; Zoued *et al*, 2016) and the crystal structure of its cytoplasmic hook-like domain has been reported from various species including EAEC, *P. aeruginosa* and *V. cholerae* (Durand *et al*, 2012; Robb *et al*, 2012; Chang & Kim, 2015; Wang *et al*, 2018)(Durand *et al*, 2012; Robb *et al*, 2012; Chang & Kim, 2015). The TssM cytoplasmic domain is comprised between the N-terminal transmembrane hairpin and a third transmembrane helix (Ma *et al*, 2009; Logger *et al*, 2016). No high-resolution structure of the TssM cytoplasmic domain is available, although a model has been built based on homology with DPY-30 and NTPases (Logger *et al*, 2016). Unfortunately, due to the poor resolution of the base, we did not succeed to confidently fit the TssL and TssM cytoplasmic domains in this density. Additional assays to improve the resolution such as the use of nanodiscs or amphipols proved to be unsuccessful (Fig EV13A). The flexibility of the TssJLM base, which did not allow for it to be resolved, might be due to the absence of other T6SS cytoplasmic components such as the baseplate. A similar observation was made for the type III secretion system, in which the presence of the cytoplasmic sorting platform orders the IM components (Hu *et al*, 2017). One may hypothesize that this flexibility is essential for the docking of the hexameric baseplate, and to accommodate the five-fold to six-fold symmetry mismatch. One alternative hypothesis is that the disorder at the centre of the base structure is caused by the presence of a lipid bilayer encircled by the TssM and TssL proteins. In the *in vivo* situation, the MC assembles first, before the recruitment of the baseplate (Durand *et al*, 2015; Brunet *et al*, 2015), and hence, one can expect that a lipid bilayer at the entrance of the TssJLM lumen would be present before baseplate docking to prevent the leakage of solutes and proton-motive force.

The high-resolution structure of the EAEC TssJLM MC also revealed new interesting and functional features. First, five additional TssJ subunits, called TssJ’, were identified in the tip complex. These TssJ’ proteins interact with the TssJ proteins of the outer pillars (TssJ.i). Mutations that interfere with TssJ’-TssJ.i interaction impair the functional integrity of the MC and hence inactivated the T6SS (Fig 4D-F). Second, we identified a 11-amino-acid loop in TssM that protrudes from each inner pillar to the centre of the channel, thus creating a constriction that is observed in the density map (Fig 3D, E). Each loop is stabilized by the adjacent loop via Asn/Gln pairings. Such weak interactions could be easily displaced by the VgrG/PAAR spike upon baseplate docking or during firing. Our mutational analyses demonstrate that this constriction is important for TssJLM MC formation and T6SS activity. Periplasmic constrictions are usual features of trans-envelope complexes. The best characterized examples include the OM T2SS and T3SS secretins, and the CsgG curli secretion channel who present one or two periplasmic gates to prevent leakage (Fig EV14A-B) (Yan *et al*, 2017; Spagnuolo *et al*, 2010; Goyal *et al*, 2014). While we do not know the role of this constriction in the T6SS, we propose that these loops may stabilize the MC in its closed conformation during the resting state. The cryo-EM structure defined two hinge regions that exhibit a certain degree of flexibility (Fig EV9B,G). These two hinges result in the formation of the two layers of pillars, with the inner layer obstructing the channel. A large conformational change of these pillars is therefore necessary to open the channel for the passage of the tube/spike complex. Interestingly, with an interaction surface of 1540 Å^2^ (∆G of 6.1 kcal/mol) the TssM inner pillars contacts are considerably less stable than the contacts within the T2SS secretin (interaction surface of 5353.7 Å^2^ and ∆G of −52.4 kcal/mol). The displacement of the pillars could be controlled by the flexibility of the hinge regions.

Based on these data, we propose a model in which the T6SS TssJLM MC is assembled in a closed state. In this conformation, five pillars are oriented toward the centre of the complex to close the complex at the outer membrane and hence to protect the cell from periplasmic leakage or from the entry of toxic compounds. This conformation is further stabilized by the interactions of the TssM protruding loops. The flexibility of the MC cytoplasmic base allows the proper positioning of the TssKFGE-VgrG baseplate, and accommodates the five-to-six symmetry. The docking of the baseplate positions the VgrG/PAAR spike in proximity to the inner membrane. Once in contact with the target cell, a signal transmitted to the baseplate triggers the contraction of the sheath, allowing the passage of the tube/spike complex through the MC. The hinge regions undergo a tectonic conformational change that opens the channel and the tip complex. The MC then returns to the resting, closing state allowing a new cycle to start (Figure 6).

**Figure 6.**
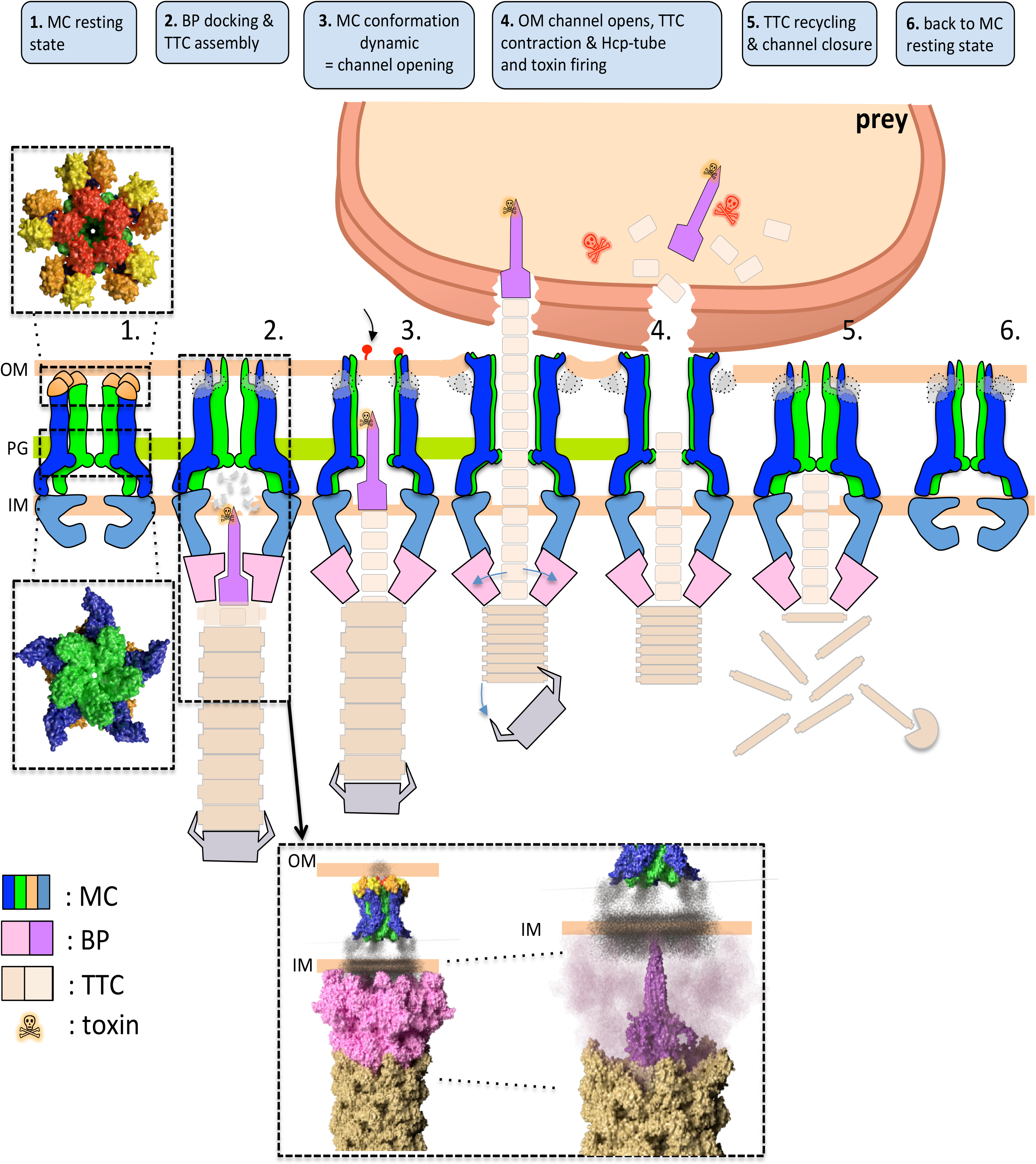
Summary of the Type 6 secretion system cycle of action. The T6SS assembly begins first with the recruitement of the membrane complex (MC) in its resting state (1). The MC recruits the baseplate (BP) and the tail tip complex (TTC) is assembled (2). The recently solved structures of the BP (No Title) and MC (this paper) in the membrane context are shown in the inset. A conformational change leads to the channel opening (3) and release of the toxin onto the VgrG spike by contraction of the TTC (4). Once the secretion has occurred, the TTC is recycled (5) and the MC can return to its resting state (6).

## Acknowledgements

We would like to thank Yoann Santin for advice on fluorescent microscopy data recording, treatment and analysis, Erney Ramírez-Aportela for help with map sharpening with LocalDeblur and Marion Decossas-Mendoza for help with grid preparation.

This work has benefitted from the facilities and expertise of the Biophysical and Structural Chemistry platform (BPCS) at IECB, CNRS UMS3033, Inserm US001, Bordeaux University, in particular we would like to thank Armel Bezault. The authors acknowledge the support and the use of resources of Instruct-ERIC and Diamond light source for the collect of the amphipoles-containing sample. ScopeM is acknowledged for instrument access at ETH Zürich.

This work was funded by the Centre National de la Recherche Scientifique, the Aix-Marseille Université, and grants from the Agence Nationale de la Recherche (ANR-14-CE14-0006-02, ANR-17-CE11-0039-01) and the Fondation pour la Recherche Médicale (DEQ20180339165) to EC. ED was supported by the INSERM and an EMBO short-term fellowship (ASTF 417 – 2015). YC is supported by a Doctoral school PhD fellowship from the FRM (ECO20160736014). VS is supported by a post-doctoral fellowship from the association Espoir contre la Mucoviscidose. LL was supported by a fourth year PhD fellowship from the FRM (FDT20160435498). MP is funded by the European Research Council, the Swiss National Science Foundation and the Helmut Horten Foundation. RF and CR were supported by IDEX Bordeaux through a “chaire d’excellence” to RF.

## Materials and methods

### Strains and media

Strains used in this study are listed in Table EV2. The E *E. coli* K-12 W3110 bearing the pUA66-*rrnB* vector (Kan^R^ and GFP^+^, (Zaslaver *et al*, 2006)) was used as recipient for antibacterial competition assays. Strains were routinely grown in lysogeny broth (LB) rich medium or in Sci-1-inducing medium (SIM; M9 minimal medium, glycerol 0.2%, vitamin B1 1 μg.mL^-1^, casaminoacids 100 mg.mL^-1^, LB 10%, supplemented or not with bactoagar 1.5%) (Brunet *et al*, 2011) with shaking at 37°C.

### Strain construction

*tssM* and *tssJ* point mutations were engineered at the native locus on the chromosome by allelic replacement using the pKO3 suicide vector (Link *et al*, 1997) into the enteroaggregative *E. coli* 17-2 strain. Briefly, 17-2 WT strain was transformed with a pKO3 plasmid in which a fragment of the *tssM* or *tssJ* gene carrying the point mutations has been cloned (see below). Insertion of the plasmid into the chromosome was selected on chloramphenicol plates at 42°C. Plasmid sequences removal was then selected on 5% sucrose plates without antibiotic and *tssM* point mutation recombinant strains were screened by PCR and confirmed by DNA sequencing (Eurofins,MWG). Chromosomal fluorescent reporter insertions into the enteroaggregative *E. coli* 17-2 strain mutated in *tssM* or *tssJ* was achieved by using a modified one-step inactivation procedure (Datsenko & Wanner, 2000) as previously described (Aschtgen *et al*, 2008b) using plasmid pKOBEG (Chaveroche *et al*, 2000). Briefly, a kanamycin cassette was amplified from plasmid pKD4 using oligonucleotide pairs carrying 5’ 50-nucleotide extensions homologous to regions adjacent to the gene to be deleted. After electroporation of 600 ng of column-purified PCR product, kanamycin-resistant clones were selected and verified by colony-PCR. The kanamycin cassette, inserted at the gene locus on the bacterial chromosome, was then excised using plasmid pCP20, leaving an FRT scars. Gene deletions were confirmed by colony-PCR and sequencing.

### Fluorescence microscopy, image treatment and analyses

Fluorescence microscopy experiments were performed as described (Brunet *et al*, 2013; Zoued *et al*, 2013). Briefly, cells were grown overnight in LB medium and diluted to *A_600nm_* ^~^ 0.04 in SIM. Exponentially growing cells (*A_600nm_* ^~^ 0.8–1) were harvested, washed in phosphate-buffered saline buffer (PBS), resuspended in PBS to *A_600nm_* ^~^ 50, spotted on a 1.5% agarose pad and covered with a cover slip. Fluorescence micrographs were captured using AxioImager M2 microscope (Zeiss) equipped with an OrcaR2 digital camera (Hamamatsu). For time lapse fluorescence microscopy, images were recorded with a Nikon Eclipse Ti microscope equipped with an Orcaflash 4.0 LT digital camera (Hamamatsu) and a perfect focus system (PFS) to automatically maintain focus so that the point of interest within a specimen is always kept in sharp focus at all times despite mechanical or thermal perturbations. Fluorescence images were acquired with a minimal exposure time to reduce bleaching and phototoxicity effects, typically 200 ms for TssB-sfGFP and 300 ms for sfGFP-TssM. For image treatment, noise and background were reduced using the ‘Subtract Background’ (20 pixels Rolling Ball) and Band plugins of imageJ (Rasband, 2012). The sfGFP foci were automatically detected using the microbeJ plugin (Ducret *et al*, 2016). Floating bars representing the number of detected foci for each strain were made using GraphPad (https://www.graphpad.com). Microscopy analyses were performed at least three times, each in technical triplicate, and a representative experiment is shown.

### Interbacterial competition assay

The antibacterial growth competition assay was performed as previously described (Flaugnatti *et al*, 2016b). Wild-type *E. coli* K-12 strain W3110 bearing the pUA66-*rrnB* plasmid (conferring kanamycin resistance and constitutive GFP fluorescence (*gfp* gene under the control of the ribosomal *rrnB* promoter, (Gueguen & Cascales, 2013) was used as recipient. Attacker and recipient cells were grown for 16 h in LB medium, diluted in SIM to allow maximal expression of the *sci-1* gene cluster (Brunet *et al*, 2011). Once the culture reached *A_600nm_*^~^ 0.8, cells were harvested and normalized to *A*_600nm_ = 0.5 in SIM. Attacker and recipient cells were mixed to a 4:1 ratio and 15-μl drops of the mixture were spotted in triplicate onto a pre-warmed dry SIM agar plate. After incubation for 4 h at 37°C, the bacterial spots were resuspended in LB and bacterial suspensions were normalized to *A*_600nm_ = 0.5. For the enumeration of viable prey cells, bacterial suspensions were serially diluted and spotted onto kanamycin LB plates. The assays were performed from at least three independent cultures, with technical triplicates and a representative technical triplicate is shown.

### Protein preparation

The expression and purification of the TssJLM complex was carried out as previously described (Durand *et al*, 2015), with the exception that the cryo-EM grids were prepared immediately after the HisTrap Elution. For the amphipole-containing sample, the Strep-Trap elution was incubated with amphipoles A8-35 (Anatrace, USA) and subjected to gel filtration on a superpose 6 (GE Healthcare, UK) to remove residual detergent.

### Cryo-EM grids preparation and data acquisition

C-flat ™ (CF-2/1-2C) grids were coated with graphene oxide as previously described (Martin *et al*, 2016). 3.5 μl of the sample at 0.2 mg.mL-1, was loaded on the copper side and then blotted on the same side for 2s in a Leica EM GP at 80% humidity and 4 °C, before being plunge frozen in liquid ethane (ࢤ184°C). Micrographs (Fig EV3B) at a nominal magnification of 120,000 X were collected in a Talos Arctica electron microscope equipped with a Falcon 3EC camera (Thermo Fisher, Waltham, MA, USA) in linear mode and with a pixel size of 1.24 Å. Dose-fractionated movie frames 20/micrograph were acquired for 1 s with a total electron flux of 120 e/Å/s. The defocus range chosen for the automatic collect was 0.7 to 2 μm, which resulted in an actual range between 0.4 to 5 μm.

For the amphipoles-containing MC collection, 3019 movies composed of 25 frames at a defocus range between 0.7 and 2μm, were collected at 1.38Å pixel size with a 5s exposure time and 15 e/pix/s exposure rate at the Krios 2 at the Diamond eBIC facility.

### Cryo-EM image processing

The 16,000 movies collected were aligned using MotionCor2, with dose weighting (6 e-/Å^2^/frame) and with 5X5 patches applied (Zheng *et al*, 2017). gCTF was used to estimate the CTF parameters (Zhang, 2016) and low quality images were discarded. Relion 2.1 (Scheres, 2012) autopicked 227,527 particles and after several rounds of 2D classification in cryosparc (Punjani *et al*, 2017) and a heterogeneous *ab initio* reconstruction (2 classes), 37,435 particles were converted using the script csparc2star.py (Asarnow, 2016) and selected for a final 2D classification in relion 2.1 (Fig EV3C), of which 36,828 particles were selected. An initial unmasked refinement using the *ab initio* model from crysoparc, gave us a resolution of 7.6 Å with 5-fold applied symmetry and a soft mask of 450 Å. This refined structure was used to do a movie refinement with all the frames and a polishing step with RELION2.1. The final masked refinement of the full structure gave a final resolution of 4.9 Å with a C5 symmetry applied, and 7.9 Å with no symmetry applied (Fig EV3E, I), The disordered tip and base were subtracted and a masked refinement around the core structure yielded a final resolution of 4.6 Å (Fig EV5B). The base focused refinement was also performed on subtracted particles, without the tip and the core regions, to a resolution of 17Å (Fig EV4F). The resolution for all densities except the base, was calculated by masked postprocessing according to the “gold standard” method using 0.143 as the FSC value cut-off, or 0.5 for the low resolution reconstruction (Rosenthal & Henderson, 2003) and the local resolution of the core was calculated by relion 2.1 (Fig EV5 C).

For figures and to build *de novo* pseudoatomic models in Coot (Emsley *et al*, 2010), the cryo-EM density was initially sharpened using phenix.autosharpen (Terwilliger *et al*, 2018) and later with LocalDeblur (Ramírez-aportela *et al*, 2018). Fitting of density, correlation calculations, molecular graphics and analyses were performed on UCSF Chimera (Pettersen *et al*, 2004). For the amphipoles dataset, 2D classes were calculated from a total of 8637 particles (Fig EV13A).

### Model building

Model building proceeded by fitting the PDB 4Y7O (Durand *et al*, 2015) into the density 2 times for each pillar, with the cross correlation being calculated using Chimera (Pettersen *et al*, 2004)(Fig EV5D). The model was manually built in Coot (Emsley *et al*, 2010) using bulky sidechains and secondary structure predictions (Kelley *et al*, 2015) as guides. The de novo structure was validated and the register adjusted using residue contact prediction (Bouvier *et al*, 2018). The map of the co-evolution contacts was aligned with those of the built PDB using the MapAlign software (Ovchinnikov *et al*, 2017). Where discrepancies were observed, the register was modified to fit the predicted contact maps (Fig EV8). The model was then refined using one round of rosetta.refine (Wang *et al*, 2016) and phenix.real_space_refine (Afonine *et al*, 2018).

### Validation of the data

The model was validated as in the protocol in Refmac5 (Murshudov *et al*, 2011). The FSC map to model was calculated with the sharpened map (FSC_sum_). The model was shaken by 0.5 Å and the FSC map to model was calculated with one Half map (FSC_work_). This refined model was then used to calculate the FSC map to model with the other Half map (FSC_free_) (Fig EV15A).

The cross correlation between each amino acid in the model and map was also calculated with phenix.real_space_refine (Afonine *et al*, 2018) (Fig EV15B) and the Molprobity score (Chen *et al*, 2010). was obtained from the online server (Table EV5) Pore radius calculations were carried out using the HOLE (Smart et al, 1996) plugin in Coot and the protein interfaces were analysed with PISA (Krissinel & Henrick, 2007).

### Strains, media and chemicals

The strains, plasmids and nucleotides used in this study are listed in Tables EV2 and 3. For the cryoET studies, *E. coli* K-12 BL21(DE3) and enteroaggregative *E. coli* EAEC strain 17-2 were used for protein overexpression before plunge freezing. Strains were routinely grown in LB-Miller or in Sci-1-inducing medium (SIM; M9 minimal medium, glycerol 0.2%, vitamin B1 1 mg ml^-1^, casaminoacids 100 mg ml^-1^, LB 10%, supplemented or not with bactoagar 1.5%) (Brunet *et al*, 2011) with shaking at 37°C. Plasmids were maintained by the addition of ampicillin (100 mg ml^-1^ for E. coli K-12, 200 mg ml^-1^ for EAEC), kanamycin (50 mg ml^-1^) or chloramphenicol (30 mg ml^-1^). Expression of genes from pRSF (in BL21) and pBAD33 (in EAEC) vectors was induced for 2-3h with 1 mM of isopropyl-b-D-thio-galactopyrannoside (IPTG) or 0.3% L-arabinose, respectively.

### Preparation of frozen-hydrated specimens

Plunge freezing was performed according to (Weiss *et al*, 2017). *E. coli* BL21 or EAEC cells were concentrated by centrifugation to an OD_600_ of 3 - 20 and then mixed with protein A – 10 nm gold conjugate (Cytodiagnostics Inc.). The higher concentrations of cells were used when preparing grids for cryo-focused ion beam (cryo-FIB) milling to form “bacterial lawns” of several layers of bacteria on top of each other. Bacterial lawns were found to be more amenable to cryo-FIB milling then individual cells. A 3 μL droplet of the sample was applied to a carbon-coated EM copper grid (R2/1, Quantifoil) that had been previously glow-discharged for 90 s at −25 mA using a Pelco easiGlow^TM^ (Ted Pella, Inc.). The grid was plunge-frozen in liquid ethane-propane (37 %/63 %) using a Mark IV Vitrobot (Thermo Fisher Scientific). The forceps were mounted in the Vitrobot (27°C, humidity 95%) and the grid was blotted from both sides or only from the backside by installing a Teflon sheet (instead of a filter paper) on the front blotting pad. Grids were stored in liquid nitrogen.

### Cryo-focused ion beam milling

Cryo-focused ion beam (cryo-FIB) milling was used to prepare samples of plunge-frozen cells that could then be imaged by electron cryotomography (Marko *et al*, 2007). Our cryo-FIB milling workflow has been detailed in (Medeiros *et al*, 2018b). Frozen grids with lawns of *E. coli* BL21 cells overexpressing TssJLM were clipped into modified Autogrids provided by J. Plitzko or a commercial prototype provided by Thermo Fisher. We then transferred the grids into the liquid nitrogen bath of a loading station (Leica Microsystems) and clamped them onto a “40° pre-tilted TEM grid holder” (Leica Microsystems). The holder with grids was shuttled from the loading station to the dual beam instrument using the VCT100 transfer system (Leica Microsystems). The holder was mounted on a custom-built cryo-stage in a Helios NanoLab600i dual beam FIB/SEM instrument (FEI). The stage temperature was maintained below −154°C during loading, milling and unloading procedures. Grid quality was checked by scanning EM (SEM) imaging (5 kV, 21 pA). The samples were then coated with a Platinum (Pt) precursor gas using the Gas Injector System. We adapted a “cold deposition” technique that was published previously (Hayles *et al*, 2007) (needle distance to target of 8 mm, temperature of the precursor gas of 27 °C, and open valve time of 5 s). Lamellae were milled in several steps. We first targeted two rectangular regions to generate a lamella with ^~^2 μm thickness with the ion beam set to 30 kV and ^~^400 pA. The current of the ion beam was then gradually reduced until the lamella reached a nominal thickness of 150-400 nm (ion beam set to ^~^25 pA). Up to 6 lamellae were milled per grid. After documentation of the lamellae by SEM imaging, the holder was brought back to the loading station using the VCT100 transfer system. The grids were unloaded and stored in liquid nitrogen.

### Electron cryomicroscopy and electron cryotomography

*E. coli* BL21 and EAEC cells (overexpressing TssJLM where indicated), cryo-FIB-processed *E. coli* BL21 cells overexpressing TssJLM, and purified TssJLM samples were examined by electron cryotomography (cryoET). Images were recorded on a Tecnai Polara TEM (Thermo Fisher Scientific) equipped with post-column GIF 2002 imaging filter and K2 Summit direct electron detector (Gatan), or on a Titan Krios TEM (Thermo Fisher Scientific) equipped with a Quantum LS imaging filter and K2 Summit (Gatan). Both microscopes were operated at 300kV and the imaging filters with a 20 eV slit width. The pixel size at the specimen level ranged from 4.93 Å to 4.05 Å. The latter pixel-sized was used for the sub-tomogram average. Tilt series covered an angular range from −60° to +60° with 2° (lamellae, sheath preparations) increments and −10 to −6 μm defocus, or in focus (0 μm defocus) when the data was collected on the Titan Krios with a Volta phase plate (Thermo Fisher Scientific) (Danev & Baumeister, 2016). The total dose of a tilt series was 60-100 e^-^/Å^2^. Tilt series and 2D projection images were acquired automatically using UCSF Tomo (Zheng *et al*, 2007) on the Tecnai Polara and SerialEM (Mastronarde, 2005) on the Titan Krios. Three-dimensional reconstructions and segmentations were generated using the IMOD program suite (Kremer *et al*, 2005).

### Sub-tomogram averaging

Tomograms used for subtomogram averaging were not CTF-corrected, as most of the particles were extracted from tomograms collected in focus with the Volta phase plate. Individual particles were identified visually in tomograms as 5-branched stars shapes in top and bottom views and as inverted-Y shapes in side views and their longitudinal axes were manually modelled with open contours in 3dmod (Kremer *et al*, 1996). The manual particle picking and first round of sub-tomogram averaging were performed with the PEET software package on tomograms that were binned by 4 (1k reconstructions). Model points, the initial motive list, and the particle rotation axes were generated using the stalkInit program from the PEET package (Nicastro *et al*, 2006). This approach allowed the definition of each structure’s longitudinal axis as the particle y-axis. 28474 individual particles extracted from cryotomograms of *E. coli* BL21 ghost and FIB-milled cells were averaged using PEET with a box size of 44 pixels in x and z, and 72 pixels in y for the final step on data binned by 2 (2k reconstruction, final pixel size 8.1 Å). A random particle was chosen as a first reference. Missing wedge compensation was activated. The final motive lists obtained after this initial average performed on tomograms that were binned by 4 were then translated and used to perform a new round of sub-tomogram averaging on tomograms that were binned by 2 (2k reconstructions). From individual particles and after analysing the resulting average, C_5_ symmetry was imposed. The Fourier shell correlation curves were calculated in PEET to estimate resolution. A cylindrical mask centred on the structure’s longitudinal axis was applied to the volumes using imodmop (IMOD package) in order to mask neighbouring structures and the membranes during averaging. 3dmod (IMOD package) and UCSF Chimera (Pettersen *et al*, 2004) were used for visualization of the averages. 3dmod was used for generating all the movies, except for the morph and the atomic model visualization in Movie 3 that were generated in UCSF Chimera.

